# Horizontal detection of post-translational modifications to an amino acid with a nanopore based on analyte volume and translocation-time

**DOI:** 10.1101/2022.09.28.507994

**Authors:** G. Sampath

## Abstract

A method is proposed for the detection of post-translational modifications (PTMs) in single amino acids (AAs) for three types of PTMs (methylation, acetylation, and phosphorylation). It is preceded by a precursor step in which the terminal residue cleaved from a peptide is identified with a set of transfer RNAs (tRNAs) in a method proposed earlier (doi: 10.36227/techrxiv.19318145.v3). The identified AA (unmodified or modified) is separated from its cognate tRNA and translocated through a nanopore under electrophoresis. The resulting current blockade level (a proxy for analyte volume) and its width (a proxy for analyte translocation time) are measured and used to identify any PTM that might be present. The theoretical volumes of the 20 proteinogenic AAs and their PTMs are computed from crystallographic data and the ratio of the volume of a modified AA to that of an unmodified one obtained. The theoretical translocation time for the 20 AAs and their PTMs through a nanopore with a bilevel voltage profile is calculated with a Fokker-Planck drift-diffusion model. A 2-D scatter plot with these two quantities is generated for each AA type. Experimentally measured blockade levels and widths for an AA, modified or unmodified, can now be compared with the AA’s scatter plot to assign a PTM for a modified AA. PTM assignment is horizontal across the PTMs for the AA because the latter has already been identified from its cognate tRNA in the precursor step, the other 19 AA types and their PTMs are not involved. Computational results are presented for 49 PTMs covering all 20 AAs and the three PTM types mentioned above.

## 1. Introduction

The study of protein structure, function, and interaction plays a major role in biological research and drug design. Cellular processes are to a considerable extent determined by protein structure, especially three-dimensional, which is determined by its amino acid sequence. As a result protein sequencing methods are at the center of analytical science research and practice [1]. The current leader in protein sequencing is mass spectrometry (MS) [2], with Edman degradation followed by spectroscopy [3] making a limited contribution. The former is based on available samples in the femto- to atto-liter range, which is somewhat limiting because only a small number of molecules of some cellular proteins may be available. As a result single molecule (SM) methods have been sought diligently over the last two decades [4]. One SM method that continues to be pursued vigorously is based on nanopores [5].

While sequencing provides primary structure information from which protein form and function can be extracted, it is also necessary to consider modifications to individual residues in a protein, which may occur in the cell before, during, or after translation from RNA to protein, and are collectively referred to as post-translational modifications (PTMs) [6]. There is increasing interest in them because many PTMs have been implicated in disease processes. As a result methods for the detection of PTMs have become a major area of investigation. Currently PTM detection is largely based on MS because of the latter’s ability to accurately separate ions produced from a protein or peptide and precisely compute mass differences among them [7]. Such detection is often aided by database techniques, which are used to assign PTMs to residues based on pattern searches over peptide sequences stored in a proteome database.

### The present work

A method to assign a PTM to an AA after the AA has been detected separately in a previous step is proposed. It is based on a combination of volumetric analysis of AAs and their PTMs and analysis of their translocation times through a nanopore. This is a SM approach that has the potential ability to assign PTMs to modified AAs in very small samples containing only a few molecules. The method described uses as its starting point a recently proposed method of AA identification [8] based on the ‘superspecificity’ [9] property of transfer RNAs (tRNAs). Thus a tRNA can only be charged with its cognate AA and not with any other. PTM identification follows AA identification, so that it is only necessary to distinguish among PTMs of the already identified AA, the other 19 AA types are not involved.

## 2. Post-translational modifications of amino acids: a brief introduction

Considering the whole proteome of an organism, PTMs are sparse in comparison. Thus only a few copies of a given protein may undergo modification. This requires methods of purification and enrichment to increase the sample size needed for analysis. A proteome may be made up of thousands of proteins, and all proteins are not equally abundant. Thus some proteins may occur in extremely small quantities in the organism, with only a few molecules being present in some cases. As a result separation may have to be done at the single cell level and detection and identification of PTMs has to be at the SM level.

A wide range of PTMs has been observed in an organism [10]; the Uniprot knowledge base [11] contains a sample annotated list. The most common ones are methylation, acetylation, phosphorylation, and glycosylation. At the bulk level PTM-specific antibodies can be used to detect specific PTMs, however antibodies are not available for every PTM. The most widely used method is MS, which can be used for detection of almost all known PTMs and is also capable of detecting unknown ones. However, as noted earlier, MS is largely a bulk-level method and requires sample sizes with 1000s to tens of 1000s of copies of a protein molecule to work with.

There is a rich and growing literature in the field of PTMs, only a small sample is referenced here. An easy-to-read introduction can be found in [12]. The use of tandem MS, one of the most successful methods available for PTM detection, is reviewed in [13,14]. Purification and enrichment strategies are covered in [15], database methods in [16], single-cell methods in [17].

Identification of AAs and their PTMs can be based on physical and chemical properties or attached tags that may be chemical or optical. Chemical identification may sometimes be based on affinity molecules that bind to a target molecule, with the resulting complex detected by methods such as staining. Detection may also be based in part on solvent properties, such as pH tuning. Tag-based methods may be optical or chemical. Physical methods are based on the measurement of a physical property such as mass, volume, charge, etc. MS is the best example of this approach. By breaking the subject molecule into ionized fragments and measuring the masses of the latter, it is able to distinguish differences down to a single atom [2], although this is often aided by pattern searches in databases [7]. The most commonly occurring PTM is phosphorylation, with close to 60% of known PTMs being phosphorylated. Other PTMs with a significant presence include acetylation, glycosylation, and methylation. The present study is limited to phosphorylation, acetylation, and methylation. A set of 49 PTMs used in the study is given in Table 1; data for the study were downloaded from the PubChem website (https://pubchem.ncbi.nlm.nih.gov).

**Table 1.** 49 PTMs distributed by AA and PTM type. The numbers in columns 2, 4, and 6 are PubChem ids.

| AA | methylated |  | acetylated |  | phosphorylated |  |
| --- | --- | --- | --- | --- | --- | --- |
| ALA | 5288725 | N-Methyl-L-alanine | 88064 | N-acetyl-L-alanine |  |  |
| CYS | 24417 | S-Methyl-L-cysteine | 12035 | N-Acetyl-L-cysteine | 3082729 | S-Phosphocysteine |
| ASP | 22880 | N-methyl-D-aspartic acid | 65065 | N-acetyl-L-aspartic acid |  |  |
| GLU | 439377 | N-Methyl-L-glutamic acid | 70914 | N-Acetyl-L-glutamic acid |  |  |
| PHE | 6951135 | N-Methyl-L-phenylalanine | 74839 | N-Acetyl-L-phenylalanine | 54382600 | N-Phosphono-L-phenylalanine |
| GLY | 1088 | N-methylglycine | 10972 | N-acetyl-glycine | 11321074 | N-phosphono-Glycine |
| HIS | 92105 | 1-Methyl-L-histidine | 75619 | Acetyl histidine | 123911 | phosphohistidine |
| ILE | 5288628 | N-Methyl-L-isoleucine | 7036275 | N-Acetyl-L-isoleucine |  |  |
| LYS | 541646 | N-methyllysine | 6991978 | acetyllysine |  |  |
| LEU | 6951123 | N-Methylleucine | 70912 | N-Acetyl-L-leucine |  |  |
| MET | 6451891 | N-Methyl-L-methionine | 448580 | N-Acetyl-L-methionine |  |  |
| ASN |  |  | 99715 | N-alpha-acetyl-L-asparagine | 54033079 | Phospho-asparagine |
| PRO |  |  | 66141 | N-acetyl-L-proline | 91440465 | Phospho-proline |
| GLN |  |  | 182230 | N-acetyl-L-glutamine | 126961722 | N-phosphono-L-glutamine |
| ARG | 4366 | Methylarginine | 67427 | N-Acetyl-L-arginine | 92150 | Phospho-L-arginine |
| SER | 7000050 | 2-methyl-L-serine | 99478 | O-acetyl-L-serine | 106 | phosphoserine |
| THR | 2724875 | O-methyl-L-threonine | 16718064 | O-Acetyl-L-Threonine | 3246323 | O-phospho-L-threonine |
| VAL | 444080 | N-Methyl-L-valine | 66789 | N-Acetyl-L-valine |  |  |
| TRP | 676159 | 1-Methyl-L-tryptophan | 700653 | N-Acetyl-L-tryptophan | 11472143 | phosphotryptophan |
| TYR | 159657 | 3-methyl-L-tyrosine | 68310 | N-Acetyl-L-tyrosine | 30819 | O-Phospho-L-tyrosine |

## 3. Analyte volume as a discriminant of AAs and their PTMs

In the present work, two physical properties are used to distinguish among an AA and its PTMs: spatial volume and the time the AA or its PTM takes in translocating through a channel. The ability to separate an AA and its PTMs from one another in the approach described here without having to consider the other 19 AAs and their PTMs is a consequence of separating AA identification from PTM identification and making it a 2-step sequential process. This latter aspect is summarized in Section 5.

### 3.1 Volumes of AAs and their PTMs

Biomolecules can be modeled as a set of spheres, one per atom, whose radii are equal to the van der Waals radii of the constituent atoms. Since the bonds between two neighboring atoms are often shorter than the van der Waals radii of the two atoms geometric biomolecule modeling involves intersecting spheres. Additionally if biomolecules are in solution (typically water) then the radius of a sphere is increased by the radius of the solvent molecule.

Geometric modeling of biomolecules is oftentimes based on techniques from the field of computational geometry [18] and uses methods such as Voronoi diagrams and/or Delaunay triangulation. As an example the volume of the union of intersecting spheres is calculated exactly with an O(n^2^) algorithm in [19].

A second approach is to use Monte Carlo simulation. This is a non-algorithmic method and can be used to compute the volume of highly irregular objects in n-dimensional space and is also quite fast. Its use in approximating the volumes of AAs and PTMs is described in detail in [20]. Coordinate data for the centers of the constituent atoms of all 20 AAs and the 49 PTMs in Table 1 were downloaded from the PubChem website. Volumes may be computed with or without taking the solvent molecule into account; van der Waals radii of different atoms are from [21].

Table 2 shows the results for the 20 proteinogenic AAs with and without the solvent molecule taken into account. The PubChem id for an AA is given in columns 2 and 7.

**Table 2.** Volumes of unmodified amino acids from Monte Carlo simulation

| AA | PubChem id | No. of atoms | Volume ($\text{\AA}^3$) | Volume ($\text{\AA}^3$) with solvent molecule | AA | PubChem id | No. of atoms | Volume ($\text{\AA}^3$) | Volume ($\text{\AA}^3$) with solvent molecule |
| --- | --- | --- | --- | --- | --- | --- | --- | --- | --- |
| ALA | 5950 | 13 | 45.69 | 50.73 | MET | 6137 | 20 | 86.21 | 104.74 |
| CYS | 5862 | 14 | 49.18 | 54.07 | ASN | 6267 | 17 | 68.15 | 79.62 |
| ASP | 5960 | 16 | 66 | 77.1 | PRO | 145742 | 17 | 64.23 | 72.26 |
| GLU | 33032 | 19 | 72.44 | 87.77 | GLN | 5961 | 20 | 82.89 | 105.89 |
| PHE | 6140 | 23 | 96.14 | 110.55 | ARG | 6322 | 26 | 111.67 | 135.16 |
| GLY | 750 | 10 | 20.66 | 21.11 | SER | 5951 | 14 | 45.21 | 49.91 |
| HIS | 6274 | 20 | 94.33 | 118.85 | THR | 6288 | 17 | 64.78 | 75.32 |
| ILE | 6306 | 22 | 69.78 | 78.35 | VAL | 6287 | 19 | 66.05 | 75.94 |
| LYS | 5962 | 24 | 92.76 | 109.4 | TRP | 6305 | 27 | 142.47 | 185.31 |
| LEU | 6106 | 22 | 77.82 | 87.79 | TYR | 6057 | 24 | 101.8 | 118.48 |

Table 3 shows the volumes of the PTMs of Table 1 with and without the solvent molecule taken into account. PubChem ids are in columns 2 and 7.

**Table 3.** Volumes of PTMs from Monte Carlo simulation

| PTM name | PubChem id | No. of atoms | Volume ($\text{\AA}^3$) | Volume ($\text{\AA}^3$) with solvent molecule | PTM name | PubChem id | No. of atoms | Volume ($\text{\AA}^3$) | Volume ($\text{\AA}^3$) with solvent molecule |
| --- | --- | --- | --- | --- | --- | --- | --- | --- | --- |
| N-Methyl-L-alanine | 5288725 | 16 | 52.14 | 59.14 | N-Acetyl-L-methionine | 448580 | 25 | 114.92 | 143.6 |
| N-acetyl-L-alanine | 88064 | 18 | 69.7 | 83.88 | N-alpha-acetyl-L-asparagine | 99715 | 22 | 88.28 | 108.72 |
| S-Methyl-L-cysteine | 24417 | 17 | 64.9 | 77.19 | Phospho-asparagine | 54033079 | 22 | 102.12 | 123.74 |
| N-Acetyl-L-cysteine | 12035 | 19 | 75.79 | 90.58 | N-acetyl-L-proline | 66141 | 22 | 94.48 | 113.73 |
| S-Phosphocysteine | 3082729 | 19 | 99.48 | 118.27 | Phospho-proline | 91440465 | 22 | 92.62 | 109.56 |
| N-methyl-D-aspartic acid | 22880 | 19 | 80.4 | 96.68 | N-acetyl-L-glutamine | 182230 | 25 | 115.82 | 148.51 |
| N-acetyl-L-aspartic acid | 65065 | 21 | 86.16 | 104.08 | N-phosphono-L-glutamine | 126961722 | 25 | 103.93 | 125.42 |
| N-Methyl-L-glutamic acid | 439377 | 22 | 97.24 | 120.73 | Methylarginine | 4366 | 29 | 119.37 | 152.4 |
| N-Acetyl-L-glutamic acid | 70914 | 24 | 111.68 | 144.13 | N-Acetyl-L-arginine | 67427 | 31 | 142.52 | 192.25 |
| N-Methyl-L-phenylalanine | 6951135 | 26 | 117.79 | 138.86 | Phospho-L-arginine | 92150 | 31 | 146.76 | 187.89 |
| N-Acetyl-L-phenylalanine | 74839 | 28 | 141.33 | 180.96 | phosphoserine | 106 | 19 | 88.01 | 105.07 |
| N-Phosphono-L-phenylalanine | 54382600 | 28 | 144.55 | 182.14 | 2-methyl-L-serine | 7000050 | 17 | 65.53 | 79.06 |
| N-methylglycine | 1088 | 13 | 28.35 | 28.85 | O-acetyl-L-serine | 99478 | 19 | 77.44 | 88.72 |
| N-acetylglycine | 10972 | 15 | 54.79 | 62.07 | O-methyl-L-threonine | 2724875 | 20 | 75.92 | 91.7 |
| N-phosphono-Glycine | 11321074 | 15 | 50.05 | 52.76 | O-Acetyl-L-Threonine | 16718064 | 22 | 95.74 | 122.86 |
| 1-Methyl-L-histidine | 92105 | 23 | 103.93 | 123.34 | O-phospho-L-threonine | 3246323 | 22 | 103.7 | 125.26 |
| Acetyl histidine | 75619 | 25 | 115.35 | 140.29 | N-Methyl-L-valine | 444080 | 22 | 87.04 | 106.93 |
| phosphohistidine | 123911 | 25 | 123.74 | 151.94 | N-Acetyl-L-valine | 66789 | 24 | 96.71 | 120.5 |
| N-Methyl-L-isoleucine | 5288628 | 25 | 109.02 | 140.2 | N-Acetyl-L-tryptophan | 700653 | 32 | 161.25 | 209.48 |
| N-Acetyl-L-isoleucine | 7036275 | 27 | 111.94 | 140.29 | 1-Methyl-L-tryptophan | 676159 | 30 | 155.73 | 203.99 |
| N-methyllysine | 541646 | 27 | 105.46 | 135.02 | phosphotryptophan | 11472143 | 32 | 182.61 | 235.95 |
| acetyllysine | 6991978 | 29 | 119.89 | 149.6 | O-Phospho-L-tyrosine | 30819 | 29 | 148.57 | 184.24 |
| N-Methylleucine | 6951123 | 25 | 97.31 | 114.99 | 3-methyl-L-tyrosine | 159657 | 27 | 118.84 | 140.1 |
| N-Acetyl-L-leucine | 70912 | 27 | 120.88 | 153.74 | N-Acetyl-L-tyrosine | 68310 | 29 | 146.67 | 187.68 |
| N-Methyl-L-methionine | 6451891 | 23 | 100.46 | 119.7 |  |  |  |  |  |

Table 4 gives the volume ratios (= volume of modified or unmodified AA/volume of unmodified AA)

**Table 4.** Volume ratios of PTMs with respect to volume of unmodified AA

| AA | Volume ratio<br>for unmodified AA | Volume ratio for PTM type<br>Methylated | Acetylated | Phosphorylated |
| --- | --- | --- | --- | --- |
| ALA | 1 | 1.17 | 1.65 |  |
| CYS | 1 | 1.43 | 1.68 | 2.18 |
| ASP | 1 | 1.25 | 1.35 |  |
| GLU | 1 | 1.38 | 1.64 |  |
| PHE | 1 | 1.26 | 1.64 | 1.65 |
| GLY | 1 | 1.37 | 2.94 | 2.5 |
| HIS | 1 | 1.04 | 1.18 | 1.28 |
| ILE | 1 | 1.79 | 1.79 |  |
| LYS | 1 | 1.23 | 1.37 |  |
| LEU | 1 | 1.31 | 1.75 |  |
| MET | 1 | 1.14 | 1.37 |  |
| ASN | 1 |  | 1.37 | 1.55 |
| PRO | 1 |  | 1.57 | 1.51 |
| GLN | 1 |  | 1.4 | 1.18 |
| ARG | 1 | 1.13 | 1.42 | 1.39 |
| SER | 1 | 1.58 | 1.78 | 2.1 |
| THR | 1 | 1.22 | 1.63 | 1.65 |
| VAL | 1 | 1.41 | 1.59 |  |
| TRP | 1 | 1.1 | 1.13 | 1.27 |
| TYR | 1 | 1.18 | 1.58 | 1.55 |

### 3.2 Covering ellipsoids of AAs and their PTMs

Another geometric approach to biomolecular modeling is based on the minimum volume ellipsoid of a set of points in 3-d space. The function *EllipsoidHull* in the R software system takes a set of points corresponding to the atom centers of a molecule and returns, among other things, the semi-axes of the smallest volume ellipsoid that encloses the set of points. Tables 5 and 6 give the results for the 20 AAs and the 49 PTMs in Table 1.

**Table 5.** Axes of minimum volume enclosing ellipsoids of the 20 proteinogenic AAs. All values are in Å (=10^−10^ m)

| AA | ALA | CYS | ASP | GLU | PHE | GLY | HIS | ILE | LYS | LEU |
| --- | --- | --- | --- | --- | --- | --- | --- | --- | --- | --- |
| semi-axis 1 | 2.56 | 2.79 | 4.21 | 6.65 | 6.29 | 2.67 | 5.43 | 5 | 12.84 | 3.85 |
| semi-axis 2 | 1.31 | 2.18 | 2.64 | 2.75 | 4.06 | 0.93 | 2.55 | 3.62 | 2.25 | 3.28 |
| semi-axis 3 | 0.86 | 0.74 | 0.75 | 0.88 | 1.16 | 0.53 | 1.78 | 1.43 | 0.95 | 1.31 |
| AA | MET | ASN | PRO | GLN | ARG | SER | THR | VAL | TRP | TYR |
| semi-axis 1 | 5.89 | 3.64 | 3.97 | 7.11 | 10.64 | 2.65 | 3.23 | 3.5 | 7.57 | 7.69 |
| semi-axis 2 | 2.74 | 2.3 | 1.75 | 2.49 | 3.14 | 2.28 | 2.78 | 2.7 | 4.71 | 3.78 |
| semi-axis 3 | 1.53 | 1.27 | 1.05 | 1.35 | 1.08 | 0.79 | 1.19 | 1.34 | 1.64 | 1.08 |

**Table 6.** Axes of minimum volume enclosing ellipsoids of the 49 PTMs in Table 1. All values are in Å (=10^−10^ m)

|  |  |  |  |  |  |  |  |  |  |  |
| --- | --- | --- | --- | --- | --- | --- | --- | --- | --- | --- |
| PubChem id | 5288725 | 88064 | 24417 | 12035 | 3082729 | 22880 | 65065 | 439377 | 70914 | 6951135 |
| semi-axis 1 | 3.1 | 5.07 | 4.44 | 5.53 | 10.02 | 5.97 | 7.07 | 10.06 | 9.05 | 9.86 |
| semi-axis 2 | 2.9 | 2.52 | 2.98 | 2.58 | 2.94 | 2.24 | 3.92 | 2.56 | 5.37 | 2.86 |
| semi-axis 3 | 0.76 | 1 | 1.06 | 1.26 | 0.82 | 0.89 | 1.01 | 0.82 | 1.18 | 1.67 |
| PubChem id | 74839 | 54382600 | 1088 | 10972 | 11321074 | 92105 | 75619 | 123911 | 5288628 | 7036275 |
| semi-axis 1 | 10.55 | 10.46 | 4.56 | 4.07 | 8.83 | 8.81 | 7.21 | 10.46 | 5.69 | 7.49 |
| semi-axis 2 | 4.84 | 4.35 | 0.87 | 1.79 | 1.09 | 2.32 | 4.41 | 2.42 | 3.54 | 3.98 |
| semi-axis 3 | 1.56 | 1.87 | 0.45 | 1.02 | 0.78 | 1.31 | 1.11 | 2.12 | 1.99 | 1.87 |
| PubChem id | 541646 | 6991978 | 6951123 | 70912 | 6451891 | 448580 | 99715 | 54033079 | 66141 | 91440465 |
| semi-axis 1 | 11.93 | 10.02 | 6.29 | 6.11 | 7.91 | 9.45 | 6.58 | 11.21 | 6.04 | 7.27 |
| semi-axis 2 | 2.86 | 3.17 | 3.37 | 4.44 | 2.99 | 4.83 | 5.36 | 2.89 | 3.43 | 3.94 |
| semi-axis 3 | 1.18 | 1.55 | 1.4 | 1.8 | 1.74 | 1.71 | 1.08 | 1.31 | 1.11 | 1.34 |
| PubChem id | 182230 | 126961722 | 4366 | 67427 | 92150 | 106 | 7000050 | 99478 | 2724875 | 16718064 |
| semi-axis 1 | 8.96 | 16.11 | 15.47 | 14.15 | 15.67 | 8.64 | 2.7 | 6.57 | 4.3 | 6 |
| semi-axis 2 | 4.59 | 2.86 | 3.4 | 5.27 | 4.97 | 2.56 | 1.65 | 2.28 | 3.42 | 3.31 |
| semi-axis 3 | 1.54 | 1.18 | 2.38 | 2.28 | 1.61 | 1.16 | 1.61 | 0.9 | 1.14 | 1.92 |
| PubChem id | 3246323 | 444080 | 66789 | 700653 | 676159 | 11472143 | 30819 | 159657 | 68310 |  |
| semi-axis 1 | 6.88 | 5.16 | 5.62 | 8.19 | 9.21 | 13.99 | 14.76 | 9.44 | 10.53 |  |
| semi-axis 2 | 3.28 | 3.32 | 4.12 | 5.83 | 5.46 | 4.57 | 3.79 | 3.98 | 4.84 |  |
| semi-axis 3 | 1.25 | 1.27 | 1.35 | 2.5 | 1.8 | 1.55 | 1.36 | 1.26 | 1.58 |  |

The data in Tables 5 and 6 are used in the next section for calculating the translocation time of an AA or PTM through a nanopore.

## 4. Translocation time through a nanopore as a discriminant of AAs and their PTMs

An electrolytic cell (e-cell) has a nanopore in a membrane that divides two chambers filled with an electrolyte and known as *cis* and *trans* [5]. With an electrical potential applied across the membrane the electrolyte is ionized, leading to an ionic current which can be measured with a detector. When an analyte is inserted into *cis* it translocates into *trans* by a combination of electrophoresis and diffusion, sometimes aided by electro-osmotic forces due to electrical charges on the wall of the pore. The intrusion of the analyte results in a decrease or blockade in the normal ionic current. The size of this blockade is roughly proportional to the volume of the analyte [22,23]. Additionally the time taken by an analyte to translocate through the pore can be used to distinguish among different analytes. Protein sequencing with nanopores is reviewed in [24]. Several studies of nanopore-based identification of PTMs are available [25-28].

Here blockade level and translocation time (which can be used to approximate dwell time) are used as discriminants of AAs and their PTMs. The measured blockade level can be compared with the theoretically calculated volume ratios in Table 4 to assign a PTM to a modified AA. The translocation time can be calculated theoretically by using a Fokker-Planck model of drift and diffusion through the pore. Following [29] (see Supplementary Information therein), E(T), the mean of the translocation time T for an analyte with diffusion coefficient D and mobility μ through a pore of length L with drift voltage V across the pore is given by

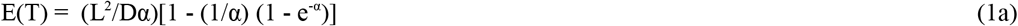

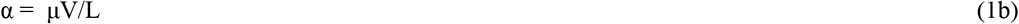

If there is no potential across the pore, or equivalently μ = 0 (uncharged analyte), translocation occurs purely through diffusion. E(T) is then given by

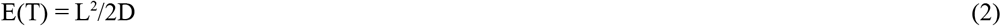

D and μ for an AA or its PTMs can be calculated from the hydrodynamic radius R_H-AA_ or R_H-PTM_ respectively. Hydrodynamic radii can be approximated from the axes of the covering ellipsoids in Tables 4 and 5. R_H_ can be approximated from the radius of a sphere R_H-sph_ with the same volume as the analyte. Thus let the analyte semi-axes be a, b, and c, where a is the major semi-axis and b and c the minor semi-axes. Then

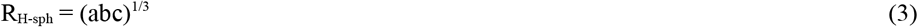

The hydrodynamic radius R_H_ for a prolate ellipsoid can be calculated with Perrin’s hydrodynamic ratio f/f_0_, which is a function of the ratio a/d, where d = √(bc) [30]. Figure 1 shows this function for prolate and oblate ellipsoids; it is given here in slightly modified form from the original in [30].

**Figure 1.**
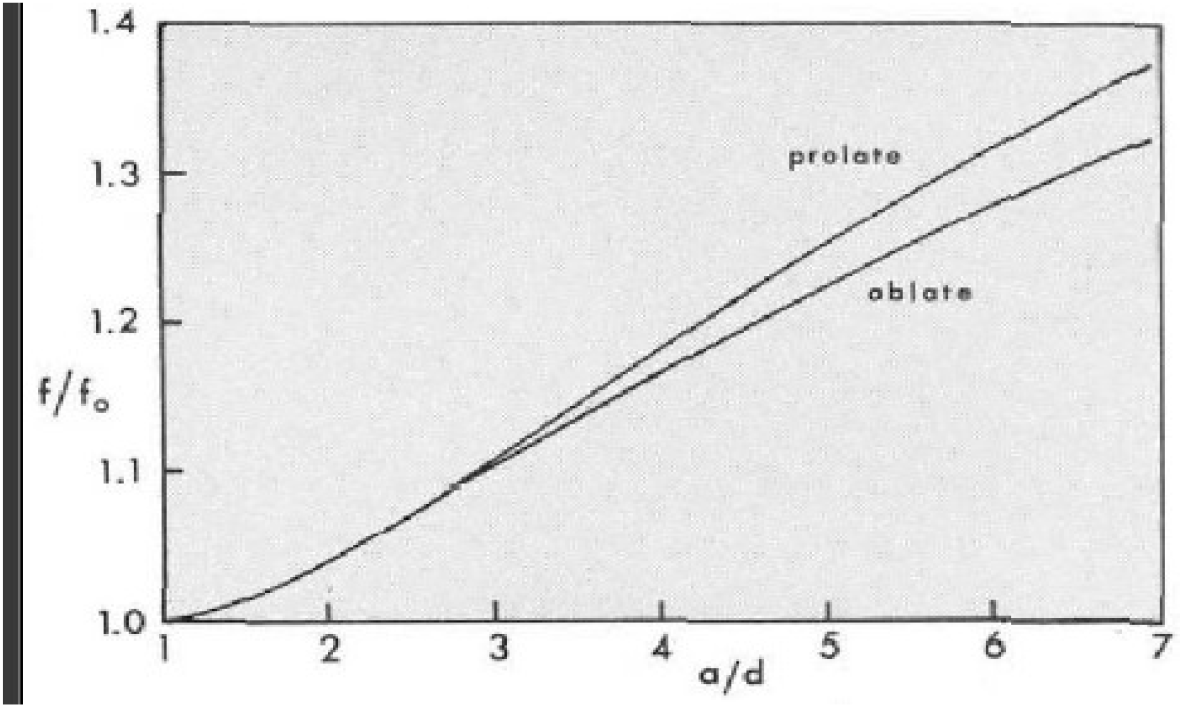
Perrin’s hydrodynamic ratio function (slightly modified version of figure in [30])

D and μ are given by

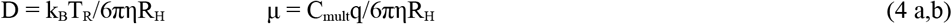

with k_B_ = Boltzmann constant (1.3806 × 10^−23^ J/K), T_R_ = room temperature (298° K), η = solvent viscosity (0.001 Pa.s for water), R_H_ = hydrodynamic radius of the analyte molecule (Å), q = electron charge (−1.619 × 10^−19^ coulomb), and C_mult_ = charge multiplier for the analyte. C_mult_ depends on the solution pH, it is calculated with the Henderson-Hasselbalch equation (see Supplement to [29]).

Tables 7 and 8 give R_H_, D, and μ for the 20 AAs and the 39 PTMs respectively.

**Table 7.** Hydrodynamic radius, mobility, and diffusion coefficient of the 20 AAs at different pH values

| AA | A | C | D | E | F | G | H | I | K | L | M | N | P | Q | R | S | T | V | W |
| --- | --- | --- | --- | --- | --- | --- | --- | --- | --- | --- | --- | --- | --- | --- | --- | --- | --- | --- | --- |
| Hydrad (Å) | 1.52 | 1.74 | 2.25 | 3.05 | 3.41 | 1.29 | 3.14 | 3.12 | 4.49 | 2.64 | 3.21 | 2.31 | 2.14 | 3.40 | 4.32 | 1.75 | 2.27 | 2.41 | 4.23 |
| pH | 11 | 10 | 7 | 7 | 12 | 12 | 12 | 12 | 7 | 12 | 12 | 12 | 12 | 12 | 7 | 12 | 12 | 12 | 12 |
| Mobility ($10^{-8}$ m<sup>2</sup>/Vs) | -1.70 | -1.93 | -1.07 | -7.54 | -6.31 | -2.41 | -6.59 | -7.12 | 4.31 | -8.05 | -7.30 | -1.05 | -1.15 | -6.57 | 4.60 | -1.49 | -1.04 | -8.99 | -4.86 |
| Diff coeff ($10^{-8}$ m<sup>2</sup>/s) | 1.43 | 1.25 | 9.70 | 7.15 | 6.40 | 1.70 | 6.95 | 7.00 | 4.87 | 8.28 | 6.81 | 9.46 | 1.02 | 6.42 | 5.05 | 1.25 | 9.61 | 9.06 | 5.16 |

**Table 8.** Hydrodynamic radius, mobility, and diffusion coefficient of the 49 PTMs in Table 1; AA properties are as in Table 7

| PTM-id | Hydrad (Å) | Diffcoeff ($10^{-10}$ m<sup>2</sup>/s) | Mobility ($10^{-8}$ m<sup>2</sup>/Vs) | PTM-id | Hydrad (Å) | Diffcoeff ($10^{-10}$ m<sup>2</sup>/s) | Mobility ($10^{-8}$ m<sup>2</sup>/Vs) |
| --- | --- | --- | --- | --- | --- | --- | --- |
| 5288725 | 1.99 | 1.10 | -1.30 | 448580 | 4.84 | 4.51 | -4.84 |
| 88064 | 2.63 | 8.30 | -9.84 | 99715 | 3.67 | 5.94 | -6.60 |
| 24417 | 2.59 | 8.42 | -1.30 | 54033079 | 4.55 | 4.79 | -5.32 |
| 12035 | 2.92 | 7.47 | -1.15 | 66141 | 3.18 | 6.87 | -7.77 |
| 3082729 | 3.89 | 5.60 | -8.62 | 91440465 | 3.79 | 5.76 | -6.52 |
| 22880 | 2.75 | 7.93 | -8.70 | 182230 | 4.54 | 4.80 | -4.92 |
| 65065 | 3.5 | 6.23 | -6.84 | 126961722 | 5.63 | 3.88 | -3.97 |
| 439377 | 3.81 | 5.74 | -6.05 | 4366 | 6.43 | 3.39 | 3.09 |
| 70914 | 4.46 | 4.89 | -5.16 | 67427 | 6.62 | 3.29 | 3.00 |
| 6951135 | 4.43 | 4.93 | -4.86 | 92150 | 6.47 | 3.37 | 3.07 |
| 74839 | 5.06 | 4.31 | -4.25 | 106 | 3.72 | 5.87 | -7.03 |
| 54382600 | 5.12 | 4.27 | -4.21 | 7000050 | 1.98 | 1.10 | -1.32 |
| 1088 | 1.19 | 1.83 | -2.60 | 99478 | 2.93 | 7.45 | -8.92 |
| 10972 | 2.17 | 1.01 | -1.43 | 2724875 | 2.69 | 8.10 | -8.78 |
| 11321074 | 3 | 7.26 | -1.03 | 16718064 | 3.59 | 6.08 | -6.59 |
| 92105 | 3.78 | 5.78 | -5.47 | 3246323 | 3.48 | 6.28 | -6.81 |
| 75619 | 3.71 | 5.88 | -5.57 | 444080 | 3 | 7.27 | -7.21 |
| 123911 | 4.65 | 4.69 | -4.44 | 66789 | 3.36 | 6.49 | -6.45 |
| 5288628 | 3.59 | 6.07 | -6.17 | 700653 | 5.17 | 4.22 | -3.98 |
| 7036275 | 4.17 | 5.23 | -5.32 | 676159 | 4.96 | 4.40 | -4.15 |
| 541646 | 4.63 | 4.72 | 4.18 | 11472143 | 5.9 | 3.70 | -3.49 |
| 6991978 | 4.5 | 4.86 | 4.31 | 30819 | 5.72 | 3.82 | -4.90 |
| 6951123 | 3.41 | 6.39 | -6.22 | 159657 | 4.36 | 5.01 | -6.43 |
| 70912 | 3.84 | 5.68 | -5.52 | 68310 | 5.07 | 4.30 | -5.53 |
| 6451891 | 3.96 | 5.51 | -5.91 |  |  |  |  |

One of the problems in nanopore sensing is the high translocation rate of analytes through the pore, which is usually beyond the bandwidth capability of the detector. A solution to this problem based on pH tuning and a bilevel voltage profile across the membrane of the e-cell is described in [29]. Figure 4b shows a three-layer membrane of the form [dielectric layer, conducting layer, dielectric layer]. By applying a positive voltage across *cis* and the middle layer and a negative voltage across the middle layer and *trans*, and adjusting the pH level to increase the mobility of an analyte, it is possible to slow down analytes enough to lower the detection bandwidth to the 1-10 Khz range. For a detailed discussion of this approach see [29].

With a multilayer membrane, the total translocation time through the stack of three pores is the sum of the translocation times through the three pores. Thus the translocation times through the two dielectric layers is calculated with Equations 1a and 1b; for the conducting layer Equation 2 is used as there is no potential gradient and transport is through pure diffusion. The mean translocation times so calculated are a lower bound, the actual times are higher because diffusion causes an analyte to randomly move back and forth when inside the pore.

The mean translocation times for the 20 AAs and 49 PTMs are given in Table 9, with volume ratios from Table 4 juxtaposed to provide a visual overview of the variations that occur in two dimensions (translocation time and volume ratio) for each AA and its PTMs

**Table 9.** Translocation times for the 49 PTMs in Table 1 through the three pore stack in Figure 4b. (Volume ratios from Table 4 juxtaposed; see main text.)

| AA | Translocation time (s) |  |  |  | Unmodified | Volume ratio |  |  |
| --- | --- | --- | --- | --- | --- | --- | --- | --- |
|  | Unmodified | AA methyl | acetyl | phosphoryl |  | AA methyl | acetyl | phosphoryl |
| A | 0.071 | 0.093 | 0.124 |  | 1 | 1.17 | 1.65 |  |
| C | 27.449 | 40.895 | 46.097 | 61.441 | 1 | 1.43 | 1.68 | 2.18 |
| D | 0.025 | 0.031 | 0.039 |  | 1 | 1.25 | 1.35 |  |
| E | 0.017 | 0.021 | 0.025 |  | 1 | 1.38 | 1.64 |  |
| F | 0.006 | 0.008 | 0.010 | 0.010 | 1 | 1.26 | 1.64 | 1.65 |
| G | 2.885 | 2.673 | 4.861 | 6.737 | 1 | 1.37 | 2.94 | 2.5 |
| H | 0.003 | 0.004 | 0.004 | 0.005 | 1 | 1.04 | 1.18 | 1.28 |
| I | 0.009 | 0.011 | 0.013 |  | 1 | 1.79 | 1.79 |  |
| K | 0.002 | 0.002 | 0.002 |  | 1 | 1.23 | 1.37 |  |
| L | 0.004 | 0.005 | 0.006 |  | 1 | 1.31 | 1.75 |  |
| M | 0.024 | 0.030 | 0.036 |  | 1 | 1.14 | 1.37 |  |
| N | 0.031 |  | 0.050 | 0.062 | 1 |  | 1.37 | 1.55 |
| P | 0.041 |  | 0.062 | 0.073 | 1 |  | 1.57 | 1.51 |
| Q | 0.011 |  | 0.015 | 0.019 | 1 |  | 1.4 | 1.18 |
| R | 0.002 | 0.004 | 0.004 | 0.004 | 1 | 1.13 | 1.42 | 1.39 |
| S | 0.099 | 0.210 | 0.112 | 0.166 | 1 | 1.58 | 1.78 | 2.1 |
| T | 0.020 | 0.024 | 0.032 | 0.031 | 1 | 1.22 | 1.63 | 1.65 |
| V | 0.005 | 0.006 | 0.007 |  | 1 | 1.41 | 1.59 |  |
| W | 0.004 | 0.005 | 0.005 | 0.006 | 1 | 1.1 | 1.13 | 1.27 |
| Y | 0.874 | 1.350 | 1.029 | 1.197 | 1 | 1.18 | 1.58 | 1.55 |

Figure 2 shows a scatter plot in these two dimensions for three AAs and their PTMs. The extent of separation seen in the plot is a measure of the ability of the proposed method to unambiguously identify and discriminate among PTMs of an AA, and, incidentally, determine if an AA is modified or not.

**Figure 2.**
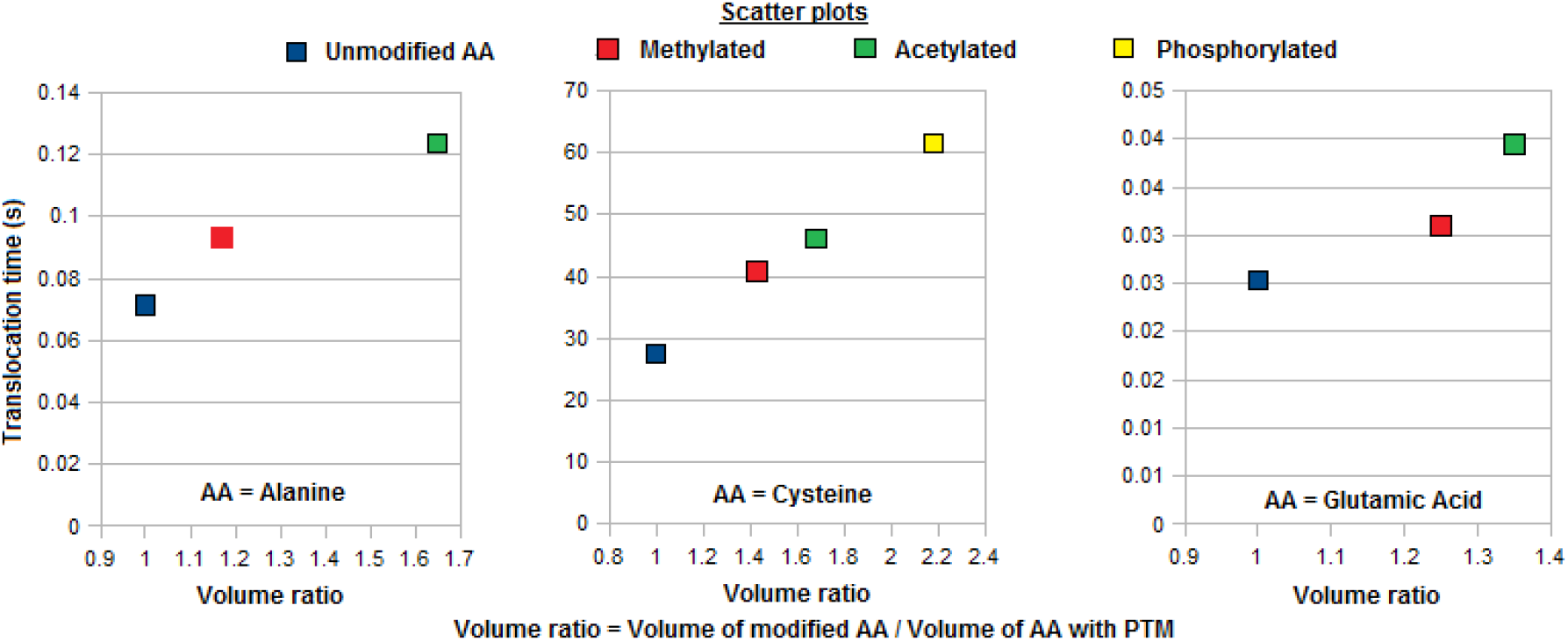
Scatter plot showing separation of three AAs and their PTMs in two dimensions

## 5. A two-step procedure for AA/PTM identification with a nanopore

AA/PTM identification/detection based on the approach given here can be implemented in two steps: 1) Identify AA (unmodified or modified); 2) Identify the PTM of the AA if any. This is a sequential process that separates AA identification from PTM identification so that the latter is restricted to selecting from the PTM candidates for the already identified AA.

In [8] a procedure is described for the unambiguous identification of free AAs cleaved from a peptide. It is based on the superspecificity property of tRNAs: a tRNA can get charged only with its cognate AA and not with any other AA. By confining each of 20 copies of an AA with a different tRNA, only one of them gets bound with its cognate tRNA. the charging event can be detected optically with TIRF (total internal fluorescence spectroscopy) if the AA is tagged with a fluorescent dye, or by translocating it through a nanopore after the bound AA is freed from the cognate tRNA. The detection is of a binary nature, so precise measurements are not needed.

The upper part of Figure 3 shows identification of AA from its cognate tRNA, the lower part shows the steps in PTM identification that follow AA identification.

**Figure 3.**
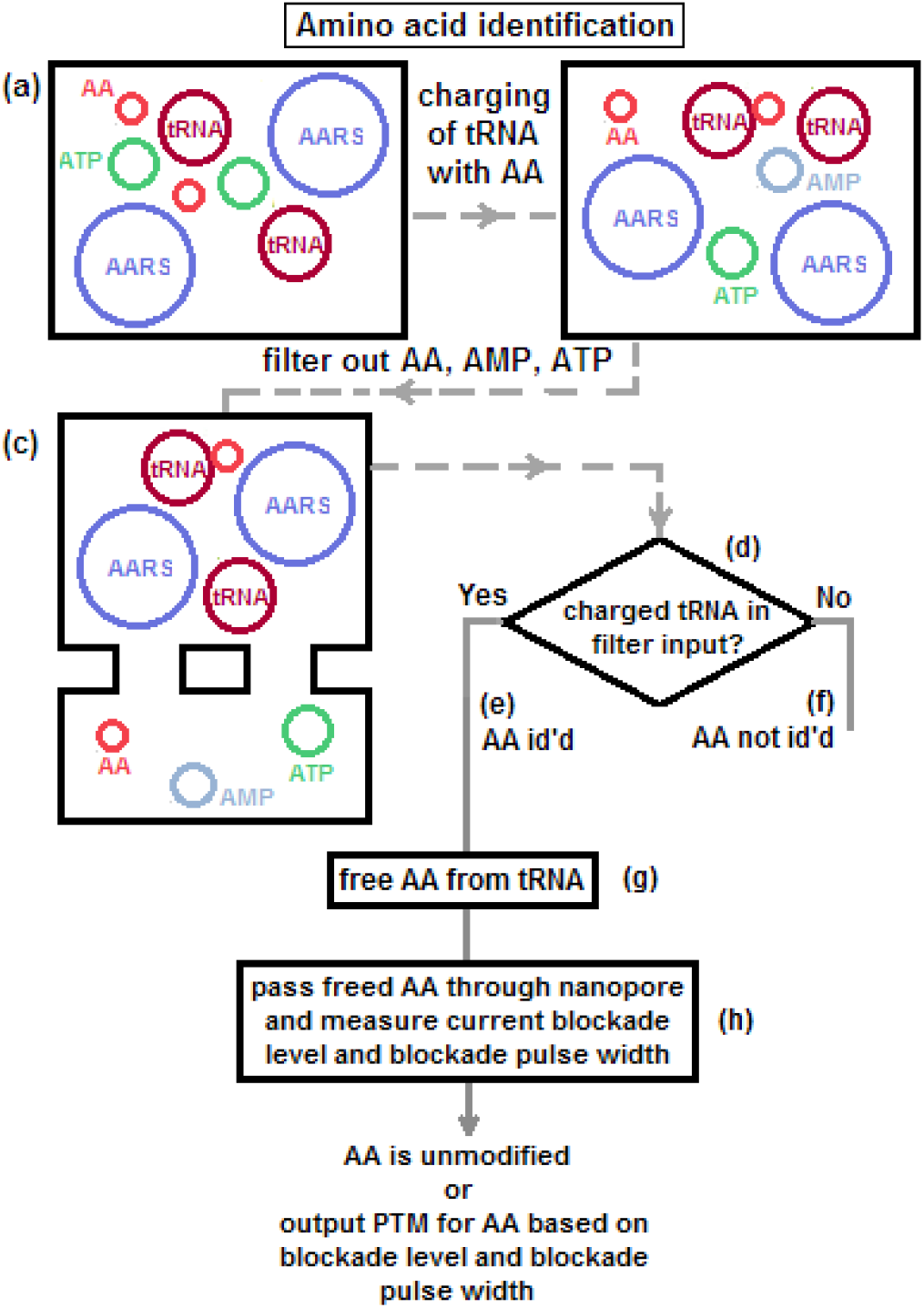
Schematic showing AA identification with tRNA followed by PTM identification with nanopore

### 4.1 AA identification with a tRNA

There are two ways to detect the AA bound to its cognate tRNA. Figure 4a shows detection of a fluorescent tag attached to the AA, Figure 4b shows detection with a nanopore after the bound AA has been deacylated and is free to translocate through a nanopore and cause the current blockade that is used to detect it. Notice that detection of the presence of AA is sufficient, identification has already implicitly occurred when tRNA gets charged with AA. (This occurs only in the identification unit with the cognate copy of AA. In the other 19 units there is no output.) For the details see [8], where a series version that can work with a single copy of AA is also outlined.

**Figure 4.**
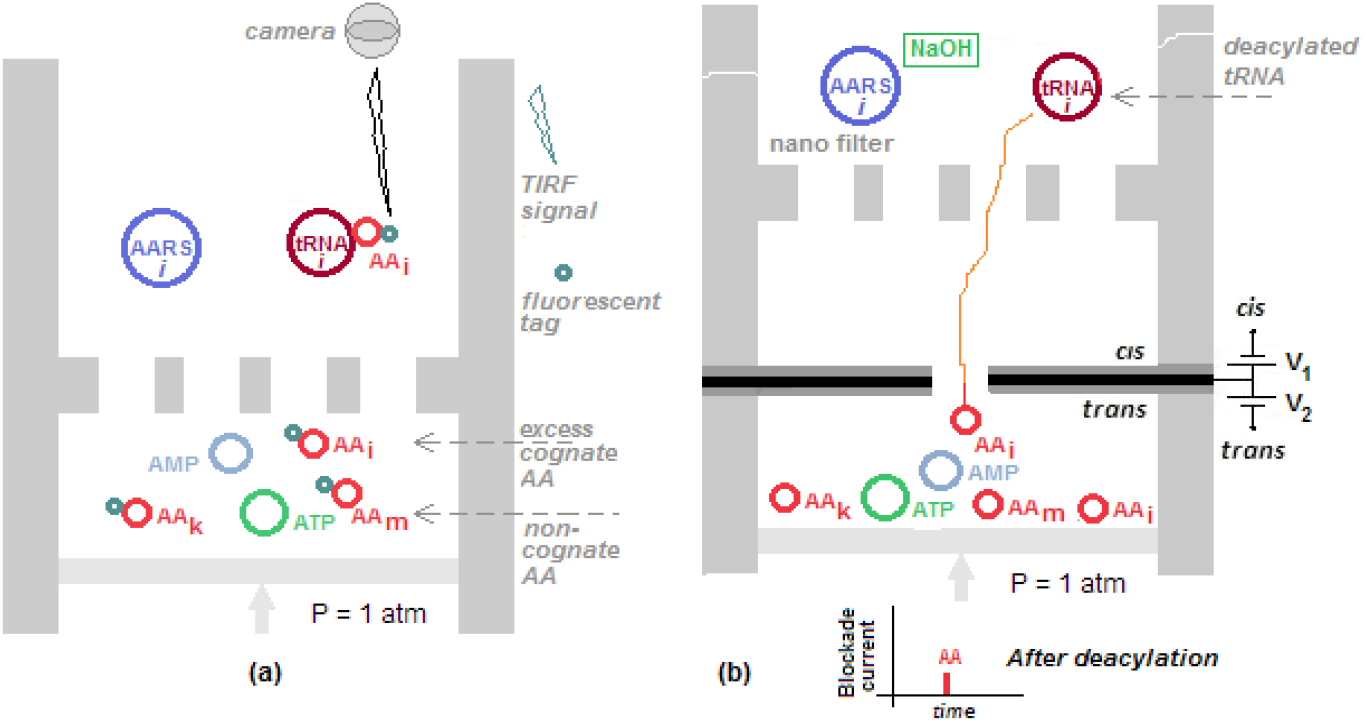
Detection of AA bound to cognate tRNA. (a) Optical detection of tag attached to AA with TIRF; (b) Electrical detection of AA after AA is freed from tRNA and passed through nanopore where occurrence of current blockade signals its presence.

### 4.2 PTM identification with a nanopore

With electrical detection, AA detection and PTM identification are combined in the nanopore. With optical detection the AA is released from the cognate tRNA and translocated through a nanopore in an e-cell and the blockade level and width measured. PTM assignment occurs horizontally across the set of PTMs for an AA without involving the other AAs. The blockade level due to a PTM is compared with the reference blockade level for the corresponding unmodified AA. The translocation time through the pore is compared with the translocation time in Table 9. The two in combination can now be used to assign a PTM to the AA if it has been modified. Equivalently, asssignment can be visually based on the scatter plot in Figure 2.

## 6. Discussion

The work reported here is an attempt to reduce the complexity of PTM detection by separating the process of identifying an amino acid in a peptide/protein from that of detecting a PTM associated with the amino acid and assigning the PTM type to it. This is possible because of the silo-like nature of the AA identification step (see discussion in [8]). This allows optimization of the characteristics of any device that comes after identification of AA has occurred through charging of a cognate tRNA. If AA identification is considered one dimension in a 2-dimensional space and PTM identification the other, then PTM identification becomes a horizontal process because only those PTMs that are associated with the identified AA are involved, the other 19 AA types and their PTMs do not play a role. If PTM identification is enhanced by database search methods then search procedures would be significantly less complex because of horizontality.

## Supporting information

AA coordinates

PTM coordinates

## Supplementary Information

Data files for 20 AAs and 49 PTMs in SDF format, slightly modified with added annotation on the first line.

## References

[1] N. Callahan, J. Tullman, Z. Kelman, and J. Marino. “Strategies for development of a next-generation protein sequencing platform”. Trends Biochem. Sci., 2019. doi:10.1016/j.tibs.2019.09.005d.

[2] E. de Hoffmann and V. Stroobant. Mass Spectrometry: Principles and Applications, 3rd edn., Wiley, 2007.

[3] R. J. Simpson. Proteins and Proteomics: A Laboratory Manual, CSHL Press, 2008.

[4] J. A. Alfaro, P. Bohländer, M. Dai, M. Filius, C. J. Howard, X. F. van Kooten, S. Ohayon, A. Pomorski, S. Schmid, A. Aksimentiev, E. V. Anslyn, G. Bedran, C. Cao, M. Chinappi, E. Coyaud, C. Dekker, G. Dittmar, N. Drachman, R. Eelkema, D. Goodlett, S. Hentz, U. Kalathiya, N. L. Kelleher, R. T. Kelly, Z. Kelman, S. H. Kim, B. Kuster, D. Rodriguez-Larrea, S. Lindsay, G. Maglia, E. M. Marcotte, J. P. Marino, C. Masselon, M. Mayer, P. Samaras, K. Sarthak, L. Sepiashvili, D. Stein, M. Wanunu, M. Wilhelm, P. Yin, A. Meller, and C. Joo. “The emerging landscape of single-molecule protein sequencing technologies”, Nature Methods 18, 604–617, 2021. doi: 10.1038/s41592-021-01143-1

[5] L. Reynaud, A. Bouchet-Spinelli, C. Raillon, and A. Buhot, “Sensing with nanopores and aptamers: a way forward”, Sensors 20, 4495, 2020.

[6] A. Holtz, N. Basisty, and B. Schilling, “Quantification and identification of post-translational modifications using modern proteomics approaches”. In: Marcus, K., Eisenacher, M., Sitek, B. (eds) Quantitative Methods in Proteomics. Methods in Molecular Biology, vol 2228. Humana, New York, NY. https://doi.org/10.1007/978-1-0716-1024-4_16

[7] M. Mann and N. L. Kelleher, “Precision proteomics: The case for high resolution and high mass accuracy”, PNAS, 105, 18137, 2008.

[8] G. Sampath, “A binary/digital approach to peptide sequencing and protein identification”. TechRxiv.org preprint, August 2022. https://doi.org/10.36227/techrxiv.19318145.v3

[9] M. Ibba and D. Söll. “The renaissance of aminoacyl-tRNA synthesis”. EMBO Reports, 2, 382–387, 2001.

[10] Anonymous. “Post-translational modifications of amino acids”, www.cellsignal.com/ptmscan.

[11] Anonymous. “Controlled vocabulary of post-translational modifications”, ptmlist.txt at www.uniprot.org.

[12] K. W. Barber and J. Rinehart, “The ABCs of PTMs”, Nat Chem Biol. 2018 February 14; 14(3): 188–192. doi:10.1038/nchembio.2572

[13] A. M. N. Silva, R. Vitorino, M. Rosário, M. Domingues, C. M. Spickett, and P. Domingues, “Post-translational modifications and mass spectrometry detection”, Free Radic. Biol. Med. 65, 925–941, 2013. doi: 10.1016/j.freeradbiomed.2013.08.184

[14] M. R. Larsen, M. B. Trelle, T. E. Thingholm, and O. N. Jensen, “Analysis of posttranslational modifications of proteins by tandem mass spectrometry: Mass spectrometry for proteomics analysis”, BioTechniques 40, 790–798, 2006. https://doi.org/10.2144/000112201

[15] S. Doll and A. L. Burlingame, “Mass spectrometry-based detection and assignment of protein post-translational modifications”, ACS Chem. Biol. 2015, 10, 63–71. DOI: 10.1021/cb500904b

[16] S. Ramazi and J. Zahiri, “Post-translational modifications in proteins: resources, tools and prediction methods”, Database, baab012, 1–20, 2021. doi:10.1093/database/baab012

[17] Y. Zhang, C. Sohn, S. Lee, H. Ahn, J. Seo, J. Cao, and L. Cai, “Detecting protein and post-translational modifications in single cells with iDentification and qUantification sEparaTion (DUET)”, Communications Biology 3, 420, 2020.

[18] M. de Berg, O. Cheong, M. van Kreveld, and M. Overmars. Computational Geometry: Algorithms and Applications. (3rd edn.), Springer-Verlag, 2008.

[19] D. Avis, B. K. Bhattacharya, and H. Imai, “Computing the volume of the union of spheres”, The Visual Computer 3, 323-328,1988.

[20] G. Sampath, “Volumetric analysis of unmodified and modified amino acids and its application to the detection of post-translational modifications (PTMs) with a nanopore”. chemrxiv preprint, September 19, 2022. doi:10.26434/chemrxiv-2022-vlmsr

[21] S. S. Batsanov, “Van der Waals radii of elements”, Inorganic Materials 37, 871–885, 2001.

[22] H. Ouldali, K. Sarthak, T. Ensslen, F. Piguet, P. Manivet, J. Pelta, J. C. Behrends, A. Aksimentiev, and A. Oukhaled, “Electrical recognition of the twenty proteinogenic amino acids using an aerolysin nanopore”, Nature Biotech., 38, 176–181, 2020. doi: 10.1038/s41587-019-0345-2

[23] X. Liu, Z. Dong, and G. Timp, “Calling the amino acid sequence of a protein/peptide from the nanospectrum produced by a sub-nanometer diameter pore”, bioRxiv.org, preprint, October 18, 2021. doi: https://doi.org/10.1101/2021.10.17.464717

[24] L. Restrepo-Pérez, C. Joo, and C. Dekker. “Paving the way to single-molecule protein sequencing”. Nature Nanotechnology 13, 786–796, 2018. https://doi.org/10.1038/s41565-018-0236-6

[25] L. Restrepo-Pérez, G. Huang, P. R. Bohländer, N. Worp, R. Eelkema, G. Maglia, C. Joo, and C. Dekker, “Resolving chemical modifications to a single amino acid within a peptide using a biological nanopore”, ACS Nano 13, 13668–13676, 2019. DOI: 10.1021/acsnano.9b05156

[26] C. B. Rosen, D. Rodriguez-Larrea, and H. Bayley, “Single-molecule site-specific detection of protein phosphorylation with a nanopore”, Nat. Biotechnol. 32, 179–181, 2014. doi: 10.1038/nbt.2799

[27] L. Restrepo-Pérez, C. H. Wong, G. Maglia, C. Dekker, and C. Joo, ““Label-free detection of post-translational modifications with a nanopore”, Nano Lett. 19, 7957–7964, 2019. DOI: 10.1021/acs.nanolett.9b03134

[28] T. Ensslen, K. Sarthak, A. Aksimentiev, and J. C. Behrends, “Resolving isomeric posttranslational modifications using a nanopore”, bioRxiv preprint, November 28, 2021. doi: 10.1101/2021.11.28.470241

[29] G. Sampath. “Slowing down an analyte in a nanopore”, biorxiv.org, February 2022. doi: 10.1101/2021.01.11.426231

[30] J. T. Edward, “Molecular volumes and the Stokes-Einstein equation”, J. Chem. Education, 47, 261–270, 1970.

